# An epigenetic score for BMI based on DNA methylation correlates with poor physical health and major disease in the Lothian Birth Cohort 1936

**DOI:** 10.1101/278234

**Authors:** Olivia KL Hamilton, Qian Zhang, Allan F McRae, Rosie M Walker, Stewart W Morris, Paul Redmond, Archie Campbell, Alison D Murray, David J Porteous, Kathryn L Evans, Andrew M McIntosh, Ian J Deary, Riccardo E Marioni

## Abstract

**Background:** The relationship between obesity and adverse health is well established, but little is known about the contribution of DNA methylation to obesity-related health outcomes. Additionally, it is of interest whether such contributions are *independent* of those attributed by the most widely used clinical measure of body mass – the Body Mass Index (BMI).

**Method:** We tested whether an epigenetic BMI score accounts for inter-individual variation in health-related, cognitive, psychosocial and lifestyle outcomes in the Lothian Birth Cohort 1936 (n=903). Weights for the epigenetic BMI score were derived using penalised regression on methylation data from unrelated Generation Scotland participants (n=2566).

**Results:** The Epigenetic BMI score was associated with variables related to poor physical health (R^2^ ranges from 0.02-0.10), metabolic syndrome (R^2^ ranges from 0.01-0.09), lower crystallised intelligence (R^2^=0.01), lower health-related quality of life (R^2^=0.02), physical inactivity (R^2^=0.02), and social deprivation (R^2^=0.02). The epigenetic BMI score (per SD) was also associated with self-reported type 2 diabetes (OR 2.25, 95 % CI 1.74, 2.94), cardiovascular disease (OR 1.44, 95 % CI 1.23, 1.69) and high blood pressure (OR 1.21, 95% CI 1.13, 1.48; all at p<0.0011 after Bonferroni correction).

**Conclusions:** Our results show that regression models with epigenetic *and* phenotypic BMI scores as predictors account for a greater proportion of all outcome variables than either predictor alone, demonstrating independent and additive effects of epigenetic and phenotypic BMI scores.

## INTRODUCTION

Body mass is a complex trait that demonstrates a high degree of variance in the general population (1). It is determined by the contribution of genetic, lifestyle, physiological and psychosocial factors, but together these factors fail to fully explain the inter-individual variation in the most widely used clinical measure of body mass, the body mass index (BMI; kg/m^2^). Through recent advances in genomic analysis we are now able to test whether obesity-related adverse health and disease may, in part, influence or be influenced by epigenetic changes, with the ultimate aim of more accurately predicting health outcomes.

Epigenetic mechanisms regulate gene expression through processes such as histone acetylation and DNA methylation (commonly the addition of a methyl group to a cytosine base in a cytosine-guanine rich sequence of DNA, or CpG), rather than changes to the DNA sequence itself. These modifications alter DNA accessibility and chromatin structure, determining whether a gene is active in a given cell at any one time (2). Several recent studies have suggested links between BMI and DNA methylation: epigenome wide association studies (EWAS) have identified CpG sites associated with BMI at loci involved in lipid and lipoprotein metabolism, blood lipid levels, and inflammatory pathways (3, 4, 5). Shah and colleagues (6) also found that methylation profiles associated with BMI accounted for 6.9% of the variance in BMI independently of genetic profiles (polygenic scores), in a group of 1,366 individuals form the Lothian Birth Cohort (LBC), which is the sample being studied in the present report.

A growing body of research examining associations between DNA methylation and health, assumes that exposures to adverse environmental stimuli induce epigenetic changes, which increase a genetically-mediated risk of disease (7). If it is the case that differential methylation profiles contribute to the risk of adverse health, there is potential utility in using epigenetic data in the prediction of health outcomes. In the current study we test whether methylation profiles associate with adverse health outcomes. We examine 1) cross-sectional associations between an epigenetic score for BMI and health-related, cognitive, psychosocial and lifestyle outcomes and 2) whether epigenetic BMI score accounts for variance of these outcome variables independently of phenotypic BMI.

## RESEARCH DESIGN AND METHODS

### The Lothian Birth Cohort 1936

The Lothian Birth Cohort 1936 (LBC1936) is a group of individuals born in 1936 who took part in the 1947 Scottish Mental Survey when they were aged 11 years (8). The survey tested the intelligence of 70,805 children in Scotland using the Moray House Test. 3,686 individuals who had taken the original test and were living in Edinburgh and the Lothians were contacted approximately 60 years later, of whom 1091 joined the LBC1936. Members of the LBC have provided a wealth of cognitive, neuropsychological, psychosocial, biological (including genomic and other ‘omics) and neuroimaging data longitudinally as part of the LBC1936 study, which is ongoing (8, 9). Recruitment and testing of the LBC1936 are described elsewhere (10).

### Ethics

Ethical permission for the LBC1936 was obtained from the Multi-Centre Research Ethics Committee for Scotland (MREC/01/0/56) and the Lothian Research Ethics Committee (LREC/2003/2/29). Written informed consent was obtained from all participants.

### DNA Methylation

DNA methylation data were assessed in whole blood samples from the LBC1936 using the Illumina HumanMethylation450 BeadChip (Illumina Inc., San Diego, CA). Full details of sample preparation and methylation typing have been reported previously (11). In brief, background correction was performed and quality control was used to remove probes with a low detection rate (P>0.01 for >5% of samples), low quality (manual inspection), low call rate (P<0.01 for <95% of probes), and samples with a poor match between genotypes and SNP control probes, with incorrectly predicted sex.

The regression weights for the LBC1936 (8) BMI epigenetic signature were derived from an independent cohort - Generation Scotland: the Scottish Family Health Study (GS; 13, 13). Full details are provided in Appendix 1.

### Phenotypic Data

Descriptive statistics for continuous outcome variables are presented in Table S1. Binary variables relating to self-reported disease history included: high blood pressure, type 1 diabetes, type 2 diabetes, high cholesterol, cardiovascular disease, leg pain, poor blood circulation, stroke, neoplasm, health issues related to the thyroid, Parkinson’s disease, arthritis and allergies. Data collection protocols for all variables are reported elsewhere (8, 14).

### Statistical Analyses

Linear regression models were used to investigate the relationship between the epigenetic BMI score and outcome variables. For each model, the R^2^ statistic represents the proportion of variance in the outcome variable that can be accounted for by the epigenetic BMI score (see model 1), phenotypic BMI score (model 2) and both epigenetic BMI and phenotypic BMI scores together (model 3). To estimate the proportion of variance accounted for by these predictors without the effect of covariates, the R^2^ statistic represents the difference between the R^2^ of the null model and the R^2^ of models 1, 2 and 3 respectively. Age at the time of testing and sex were included as covariates in all models. Height was included as an additional covariate in models for time taken to walk 6 metres, forced expiratory volume (lung function), and general physical health. The models are as follows:

***Null model:*** *variable of interest ∼ age at time of testing + sex*

***Model 1:*** *variable of interest ∼ epigenetic BMI score + age at time of testing + sex*

***Model 2:*** *variable of interest ∼ phenotypic BMI + age at time of testing + sex*

***Model 3:*** *variable of interest ∼ epigenetic BMI score + phenotypic BMI + age at time of testing + sex*

Logistic regressions were conducted for self-reported disease history variables as these had binary outcomes (disease/no disease). Similar to the models above, logistic regressions were carried out for each variable to test whether epigenetic BMI, phenotypic BMI, and an additive model combining epigenetic BMI score and phenotypic BMI, were associated with disease history. For all analyses a Bonferroni corrected level of significance was used (p<0.05/n_tests=0.0011). Analyses were carried out in R version 3.4.0 (15).

### Sensitivity analysis

For variables that associated with epigenetic BMI, we carried out further sensitivity analyses, by including a polygenic score for BMI (for details see Appendix 2) as a predictor in the additive model with epigenetic BMI score and phenotypic BMI (model 4).

***Model 4:*** *variable of interest ∼ epigenetic BMI score + phenotypic BMI + genetic BMI score + age at time of testing + sex*

## RESULTS

This study included 903 individuals (female n=446, 49.4%), with a mean age of 69.5 years.

### Epigenetic BMI score is associated with outcomes related to physical health, biomarkers of metabolic syndrome, crystallised intelligence and social deprivation

Linear regressions were carried out to test whether the epigenetic score for BMI generated in GS was associated with health-related, cognitive, psychosocial, and lifestyle outcomes in the LBC1936. Only associations with a p value <0.0011 (i.e. after Bonferroni correction) are presented here - full results are presented in Table 1.

**Table 1:** Linear regressions of epigenetic BMI score, phenotypic BMI and epigenetic + phenotypic BMI on LBC1936 outcome variables.

|  | Epigenetic BMI |  |  |  | Phenotypic BMI |  |  |  | Additive model |  |  |  |  |  |
| --- | --- | --- | --- | --- | --- | --- | --- | --- | --- | --- | --- | --- | --- | --- |
|  | n | standardised<br>beta (SE) | raw p | R<sup>2</sup> | n | standardised<br>beta (SE) | raw p | R<sup>2</sup> | n | standardised<br>beta (SE) | raw p | standardised<br>beta (SE) | raw p | R<sup>2</sup> |
| **Physical Health** |  |  |  |  |  |  |  |  |  |  |  |  |  |  |
| Height (cm) | 901 | -0.050 (0.02) | 0.030 | 0.00 | 902 | -0.072 (0.02) | 0.002 | 0.00 | 900 | -0.029 (0.02) | 0.228 | -0.063 (0.02) | 0.009 | 0.31 |
| Weight (kg) | 900 | 0.236 (0.03) | **5.5x10<sup>-16</sup>** | 0.06 | 902 | 0.786 (0.01) | **<2x10<sup>-16</sup>** | 0.61 | 900 | -0.013 (0.02) | 0.361 | 0.790 (0.01) | **<2x10<sup>-16</sup>** | 0.61 |
| BMI (kg/m<sup>2</sup>) | 900 | 0.315 (0.03) | **<2x10<sup>-16</sup>** | 0.10 |  |  |  |  |  |  |  |  |  |  |
| Time to walk six metres (sec) | 897 | 0.140 (0.02) | **1.7x10<sup>-05</sup>** | 0.02 | 898 | 0.264 (0.03) | **2.3x10<sup>-16</sup>** | 0.07 | 896 | 0.064 (0.03) | 0.052 | 0.242 (0.03) | 5.31E-13 | 0.07 |
| Right hand grip strength (kg) | 897 | -0.027 (0.02) | 0.211 | 0.00 | 898 | -0.036 (0.02) | 0.087 | 0.00 | 896 | -0.015 (0.02) | 0.506 | -0.032 (0.02) | 0.153 | 0.00 |
| Forced expiratory volume in 1 sec<br>(litres) | 899 | -0.131 (0.02) | **6.3x10<sup>-08</sup>** | 0.02 | 900 | -0.061 (0.02) | 0.013 | 0.00 | 898 | -0.124 (0.03) | **1.2x10<sup>-06</sup>** | **-0.022 (0.03)** | **0.387** | 0.02 |
| General physical health* | 891 | -0.130 (0.02) | **4x10<sup>-08</sup>** | 0.02 | 892 | -0.155 (0.02) | **4.5x10<sup>-11</sup>** | 0.02 | 890 | -0.088 (0.02) | **3.2x10<sup>-04</sup>** | **-0.127 (0.02)** | **2.6x10<sup>-07</sup>** | 0.03 |
| **Mental Health** |  |  |  |  |  |  |  |  |  |  |  |  |  |  |
| HADS anxiety subscale† | 900 | -0.038 (0.03) | 0.244 | 0.00 | 901 | -0.046 (0.03) | 0.159 | 0.00 | 899 | -0.028 (0.04) | 0.420 | -0.036 (0.03) | 0.296 | 0.00 |
| HADS depression subscale | 897 | 0.073 (0.03) | 0.029 | 0.01 | 898 | 0.129 (0.03) | **1.2x10<sup>-04</sup>** | 0.02 | 896 | 0.035 (0.04) | 0.318 | 0.117 (0.04) | **0.001** | 0.02 |
| HADS total score | 897 | 0.007 (0.03) | 0.841 | 0.00 | 898 | 0.032 (0.03) | 0.344 | 0.00 | 896 | -0.005 (0.04) | 0.879 | 0.034 (0.04) | 0.334 | 0.00 |
| **Cognitive** |  |  |  |  |  |  |  |  |  |  |  |  |  |  |
| WAIS Symbol search‡ | 897 | -0.086 (0.03) | 0.008 | 0.01 | 898 | -0.073 (0.03) | 0.03 | 0.01 | 896 | -0.070 (0.03) | 0.040 | -0.050 (0.03) | 0.142 | 0.01 |
| WAIS Digit symbol | 896 | -0.089 (0.03) | 0.006 | 0.01 | 897 | -0.121 (0.03) | **2.0x10<sup>-04</sup>** | 0.01 | 895 | -0.057 (0.03) | 0.094 | -0.101 (0.03) | 0.003 | 0.02 |
| WAIS Matrix reasoning | 897 | -0.066 (0.03) | 0.045 | 0.00 | 898 | -0.125 (0.03) | **1.5x10<sup>-04</sup>** | 0.02 | 896 | -0.029 (0.04) | 0.407 | -0.117 (0.03) | **0.001** | 0.02 |
| WAIS Letter number sequencing | 891 | -0.067 (0.03) | 0.041 | 0.00 | 892 | -0.126 (0.03) | **1.3x10<sup>-04</sup>** | 0.02 | 890 | -0.030 (0.03) | 0.377 | -0.116 (0.03) | **0.001** | 0.02 |
| WAIS Backwards digit span | 900 | -0.038 (0.03) | 0.250 | 0.00 | 901 | -0.120 (0.03) | **2.6x10<sup>-04</sup>** | 0.01 | 899 | 0.001 (0.04) | 0.981 | -0.121 (0.03) | **0.001** | 0.01 |
| WAIS Block design | 897 | -0.072 (0.03) | 0.028 | 0.00 | 898 | -0.038 (0.03) | 0.24 | 0.00 | 896 | -0.065 (0.04) | 0.061 | -0.017 (0.03) | 0.614 | 0.01 |
| General fluid intelligence§ | 885 | -0.095(0.03) | 0.004 | 0.01 | 886 | -0.141 (0.03) | **1.4x10<sup>-05</sup>** | 0.02 | 884 | -0.055 (0.03) | 0.104 | -0.123 (0.03) | **3.1x10<sup>-04</sup>** | 0.02 |
| National Adult Reading Test | 899 | -0.118 (0.03) | **3x10<sup>-04</sup>** | 0.01 | 900 | -0.178 (0.03) | **4.7x10<sup>-08</sup>** | 0.03 | 898 | -0.067 (0.03) | 0.048 | -0.157 (0.03) | **4.3x10<sup>-06</sup>** | 0.04 |

Table 1 continued.
|  |  |  |  |  |  |  |  |  | Additive model |  |  |  |  |  |
| --- | --- | --- | --- | --- | --- | --- | --- | --- | --- | --- | --- | --- | --- | --- |
|  |  |  |  |  |  |  |  |  | Epigenetic BMI |  |  | Phenotypic BMI |  |  |
|  | n | standardised<br>beta (SE) | raw p | R<sup>2</sup> | n | standardised<br>beta (SE) | raw p | R<sup>2</sup> | n | standardised<br>beta (SE) | raw p | standardised<br>beta (SE) | raw p | R<sup>2</sup> |
| **Psychosocial factors** |  |  |  |  |  |  |  |  |  |  |  |  |  |  |
| IPIP conscientiousness\|\| | 782 | -0.069 (0.04) | 0.054 | 0.01 | 783 | -0.168 (0.04) | **3.9x10<sup>-06</sup>** | 0.03 | 781 | -0.016 (0.04) | 0.659 | -0.164 (0.04) | **1.2x10<sup>-05</sup>** | 0.03 |
| IPIP extraversion | 784 | -0.002 (0.04) | 0.965 | 0.00 | 785 | 0.028 (0.04) | 0.452 | 0.00 | 783 | -0.011 (0.04) | 0.765 | 0.031 (0.04) | 0.429 | 0.00 |
| IPIP agreeableness | 783 | -0.052 (0.03) | 0.122 | 0.00 | 784 | -0.065 (0.03) | 0.057 | 0.00 | 782 | -0.034 (0.04) | 0.332 | -0.054 (0.04) | 0.129 | 0.00 |
| IPIP emotional stability | 781 | -0.006 (0.04) | 0.877 | 0.00 | 782 | -0.041 (0.04) | 0.267 | 0.00 | 780 | 0.008 (0.04) | 0.831 | -0.045 (0.04) | 0.250 | 0.00 |
| IPIP intellect/imagination | 780 | -0.079 (0.04) | 0.027 | 0.01 | 781 | -0.078 (0.04) | 0.034 | 0.01 | 779 | -0.061 (0.04) | 0.107 | -0.059 (0.04) | 0.132 | 0.01 |
| WHOQOL physical health¶ | 792 | -0.144 (0.04) | **5.1x10<sup>-5</sup>** | 0.02 | 793 | -0.276 (0.04) | **1.5x10<sup>-14</sup>** | 0.07 | 791 | -0.064 (0.04) | 0.078 | -0.256 (0.04) | **1.1x10<sup>-11</sup>** | 0.08 |
| WHOQOL psychological. health | 793 | -0.042 (0.04) | 0.243 | 0.00 | 794 | -0.110 (0.04) | 0.002 | 0.01 | 792 | -0.008 (0.04) | 0.835 | -0.108 (0.04) | 0.005 | 0.01 |
| WHOQOL social relationships | 793 | -0.045 (0.04) | 0.211 | 0.00 | 794 | -0.047 (0.04) | 0.195 | 0.00 | 792 | -0.032 (0.04) | 0.399 | -0.037 (0.04) | 0.330 | 0.00 |
| WHOQOL environment | 792 | -0.079 (0.04) | 0.027 | 0.01 | 793 | -0.122 (0.04) | 0.001 | 0.01 | 791 | -0.044 (0.04) | 0.241 | -0.109 (0.04) | 0.005 | 0.02 |
| **Bloods and Biomarkers** |  |  |  |  |  |  |  |  |  |  |  |  |  |  |
| HBA1C (%total) | 898 | 0.252 (0.03) | **2x10<sup>-14</sup>** | 0.06 | 899 | 0.240 (0.03) | **3.1x10<sup>-13</sup>** | 0.06 | 897 | 0.196 (0.03) | **8.6x10<sup>-09</sup>** | **0.179 (0.03)** | **1.3x10<sup>-07</sup>** | 0.09 |
| Fibrinogen (g/L) | 886 | 0.107 (0.03) | 0.001 | 0.01 | 887 | 0.101 (0.03) | 0.003 | 0.01 | 885 | 0.084 (0.04) | 0.017 | 0.074 (0.04) | 0.037 | 0.02 |
| Blood derived C reactive protein<br>(mg/L) | 891 | 0.114 (0.03) | **5.9x10<sup>-4</sup>** | 0.01 | 892 | 0.181 (0.03) | **4.6x10<sup>-08</sup>** | 0.03 | 890 | 0.063 (0.04) | 0.069 | 0.161 (0.03) | **4x10<sup>-06</sup>** | 0.04 |
| Diastolic blood pressure standing | 480 | 0.083 (0.05) | 0.087 | 0.01 | 481 | 0.273 (0.05) | **2.7x10<sup>-08</sup>** | 0.06 | 479 | 0.007 (0.05) | 0.886 | 0.270 (0.05) | **1.4x10<sup>-07</sup>** | 0.06 |
| Systolic blood pressure sitting | 716 | 0.055 (0.04) | 0.145 | 0.00 | 717 | 0.142 (0.04) | **2.4x10<sup>-04</sup>** | 0.02 | 715 | 0.016 (0.04) | 0.677 | 0.137 (0.04) | 0.001 | 0.02 |
| Diastolic blood pressure standing | 716 | 0.035 (0.04) | 0.355 | 0.00 | 717 | 0.246 (0.04) | **1.4x10<sup>-10</sup>** | 0.06 | 715 | -0.037 (0.04) | 0.336 | 0.257 (0.04) | **1.5x10<sup>-10</sup>** | 0.06 |
| Systolic blood pressure standing | 716 | 0.107 (0.04) | 0.005 | 0.01 | 717 | 0.268 (0.04) | **1.9x10<sup>-12</sup>** | 0.07 | 715 | 0.033 (0.04) | 0.386 | 0.258 (0.04) | **8.6x10<sup>-11</sup>** | 0.07 |
| Triglyceride (mmol/l) | 815 | 0.311 (0.03) | **<2x10<sup>-16</sup>** | 0.09 | 816 | 0.276 (0.03) | **7.1x10<sup>-16</sup>** | 0.08 | 814 | 0.248 (0.04) | **2.4x10<sup>-12</sup>** | **0.203 (0.03)** | **4.0x10<sup>-09</sup>** | 0.13 |
| Total cholesterol (mmol/l) | 890 | -0.093 (0.03) | 0.004 | 0.01 | 892 | -0.057 (0.03) | 0.073 | 0.01 | 889 | -0.081 (0.03) | 0.016 | -0.031 (0.03) | 0.359 | 0.01 |
| High-Density Lipoprotein<br>Cholesterol (HDL; mmol/l) | 818 | -0.289 (0.03) | **<2x10<sup>-16</sup>** | 0.08 | 819 | -0.277 (0.03) | **< 2x10<sup>-16</sup>** | 0.08 | 817 | -0.223 (0.03) | **3.1x10<sup>-11</sup>** | **-0.211 (0.03)** | **1.6x10<sup>-10</sup>** | 0.12 |
| HDL Ratio (HDL<br>cholesterol/total cholesterol) | 815 | 0.185 (0.04) | **1.6x10<sup>-07</sup>** | 0.03 | 816 | 0.185 (0.03) | **9.4x10<sup>-08</sup>** | 0.03 | 814 | 0.141 (0.04) | **1.1x10<sup>-04</sup>** | **0.144 (0.04)** | **6.1x10<sup>-05</sup>** | 0.05 |

**Table 1 continued.**
|  | Epigenetic BMI |  |  |  | Phenotypic BMI |  |  |  | Additive model |  |  |  |  |  |
| --- | --- | --- | --- | --- | --- | --- | --- | --- | --- | --- | --- | --- | --- | --- |
|  | n | standardised<br>beta (SE) | raw p | R<sup>2</sup> | n | standardised<br>beta (SE) | raw p | R<sup>2</sup> | n | standardised<br>beta (SE) | raw p | standardised<br>beta (SE) | raw p | R<sup>2</sup> |
| **Lifestyle** |  |  |  |  |  |  |  |  |  |  |  |  |  |  |
| Scottish Index of Multiple Deprivation | 893 | -0.135 | **3.9x10<sup>-05</sup>** | **0.02** | 894 | -0.120 | **2.4x10<sup>-04</sup>** | **0.02** | 892 | -0.107 | 0.002 | -0.087 (0.03) | 0.011 | 0.03 |
| Units alcohol per week | 901 | -0.029 | 0.365 | 0.00 | 902 | -0.026 | 0.425 | 0.00 | 900 | -0.021 | 0.528 | -0.020 (0.03) | 0.565 | 0.00 |
| Self-reported physical activity# | 792 | -0.134 | **1.5x10<sup>-04</sup>** | **0.02** | 793 | -0.184 | **3.6x10<sup>-07</sup>** | **0.03** | 791 | -0.086 | 0.019 | -0.155 (0.04) | **4.2x10<sup>-05</sup>** | 0.04 |
| Mediterranean diet ** | 719 | -0.095 | 0.011 | 0.01 | 720 | -0.019 | 0.633 | 0.00 | 718 | -0.098 | 0.012 | 0.011 (0.04) | 0.779 | 0.01 |
| Health aware diet | 719 | -0.047 | 0.190 | 0.00 | 721 | -0.062 | 0.092 | 0.00 | 718 | -0.030 | 0.416 | -0.052 (0.04) | 0.183 | 0.01 |
| Conventional diet | 719 | 0.076 | 0.040 | 0.00 | 720 | 0.114 | 0.003 | 0.01 | 718 | 0.045 | 0.243 | 0.100 (0.04) | 0.012 | 0.01 |
| Sweet foods | 719 | 0.061 | 0.107 | 0.00 | 720 | -0.081 | 0.041 | 0.01 | 718 | 0.092 | 0.021 | -0.109 (0.04) | 0.008 | 0.01 |

\* Principal component analysis using grip strength, forced expiratory volume and six metre walk time
† Hospital Anxiety and Depression Score
‡ Weschler Adult Intelligence Scale
§ Scores from the first unrotated component of a principal components analysis of letter number sequence, matrix reasoning, backwards digit span, block design, digit symbol & symbol search were extracted and labelled general physical health
|| International Personality Item Pool personality subtypes
¶ World Health Organisation Quality of Life

Epigenetic BMI was strongly associated with phenotypic BMI (p<2 × 10^−16^) and accounted for 10% of its variation. Increases in epigenetic BMI score also correlated with poorer performance on all other physical health variables, but only correlations with 6 metre walk time, lung function and general physical health remained after correction for multiple comparisons. As epigenetic BMI score increased by 1 SD, time taken to walk 6 metres increased by 0.1 seconds (p=1.7 × 10^−05^, R^2^= 0.02), forced expiratory volume decreased by 0.1 litres (p=6.3 × 10^−08^, R^2^=0.02) and general physical health decreased by 0.13 SDs (p=4 × 10^−08^, R^2^=0.02). Epigenetic BMI score demonstrated negative correlations with blood biomarker variables: HbA1c (p=2 × 10^−14^, R^2^= 0.06), C-reactive protein (p=5.9 × 10^−4^, R^2^= 0.01), triglycerides (p<2 × 10^−16^, R^2^= 0.09), high density lipoprotein cholesterol (HDL; p<2 × 10^−16^, R^2^=0.08) and HDL ratio (HDL cholesterol/total cholesterol; p**=**1.6 × 10^−07^, R^2^=0.03). Finally, epigenetic BMI score was negatively associated with crystallised intelligence as measured by the National Adult Reading Test (NART; p=3 × 10^−04^, R^2^=0.01), Scottish Index of Multiple Deprivation score (p=3.9 × 10^−05^, R^2^= 0.02), physical health-related quality of life (p=5.1 × 10^−5^, R^2^= 0.02) and self-reported physical activity (p=1.5 × 10^−04^, R^2^= 0.02)

## Epigenetic BMI score and phenotypic BMI account for independent proportions of variance

Phenotypic BMI was associated with the same variables as epigenetic BMI score: time to walk 6 metres (p=2.3 × 10^−16^, R^2^= 0.07), general physical health (p=4.5 × 10^−11^, R^2^=0.02), HbA1c (p=3.1 × 10^−13^, R^2^= 0.06), C-reactive protein (p=4.6 × 10^−08^, R^2^= 0.03), triglycerides (p=7.1 × 10^−16^, R^2^= 0.08), HDL cholesterol (p**<**2 × 10^−16^, R^2^=0.08), HDL ratio (p=9.4 × 10^−08^, R^2^=0.03), crystallised intelligence (p=4.7 × 10^−08^, R^2^=0.03), the Scottish Index of Multiple Deprivation (p=2.4 × 10^−04^, R^2^= 0.02), physical health-related quality of life (p=1.5 × 10^−14^, R^2^= 0.07) and self-reported level of physical activity (p=3.6 × 10^−07^, R^2^= 0.03). With the exception of lung function and triglycerides, phenotypic BMI accounted for a greater amount of variance of all outcome variables, relative to epigenetic BMI,

Variables that associated with epigenetic BMI were included in additive linear regression models, with both epigenetic and phenotypic BMI score as predictors. These models showed that epigenetic and phenotypic BMI explain partially independent proportions of the variance. This pattern was most striking for biomarker variables related to metabolic syndrome – additive models that demonstrated associations with these biomarkers are presented in figure 1.

**Figure 1:**
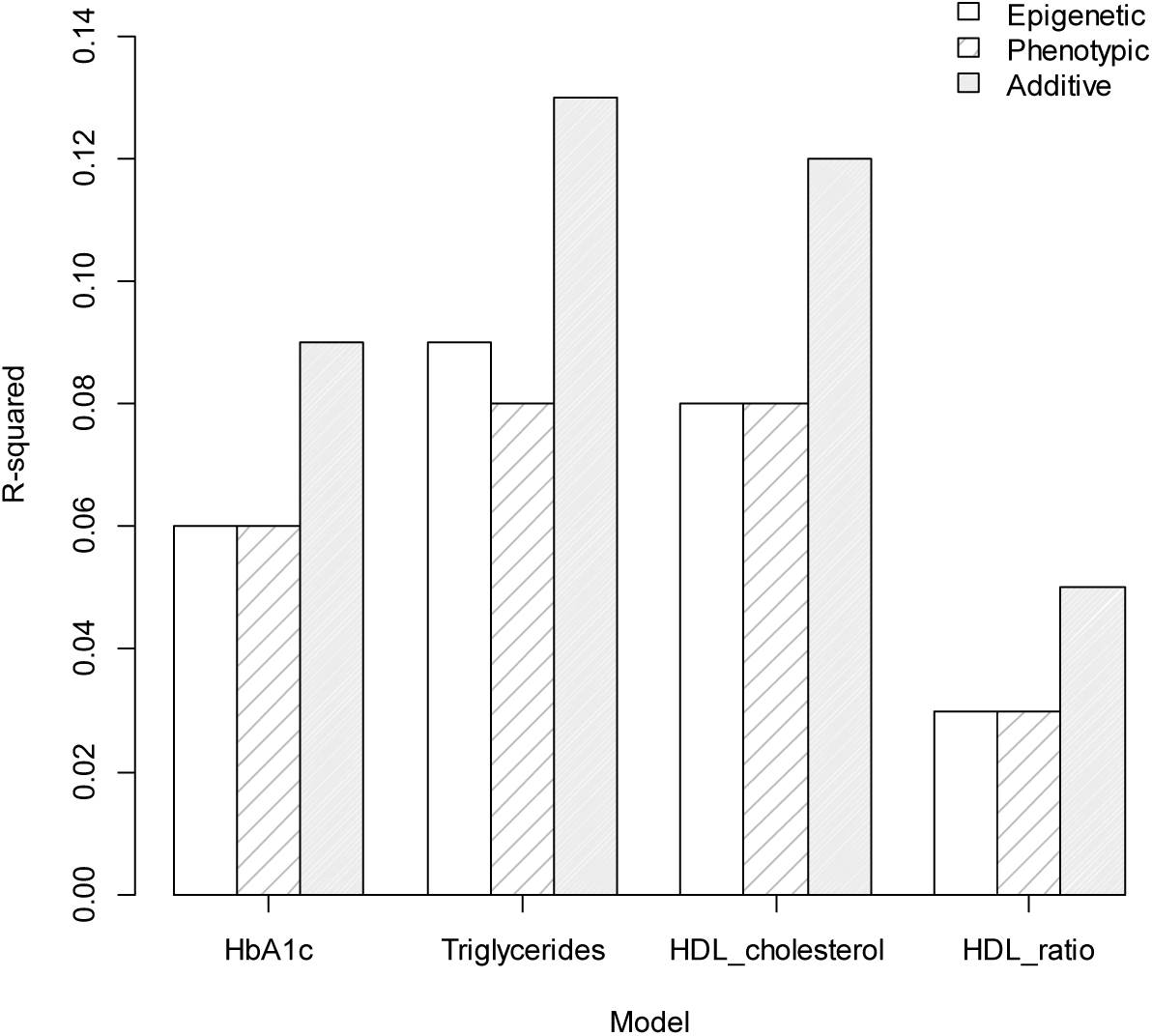
Plot of R^2^ statistics from linear regression analyses in which epigenetic BMI, phenotypic BMI and epigenetic + phenotypic BMI respectively associated with biomarker variables at p> 0.0011. The plots demonstrate the general pattern observed, with the additive model accounting for a greater proportion of variation than models including epigenetic BMI score only, or phenotypic BMI only.

## A polygenic score for BMI did not increase R ^2^ of additive models

Tables S2 and S3 present results of linear and logistic regression models in which a polygenic score for BMI was included as a predictor variable alongside epigenetic BMI score and phenotypic BMI. In these additive models, a polygenic BMI score did not associate with any of the variables tested.

A linear regression model with both epigenetic BMI score and polygenic BMI score, polygenic BMI showed a strong association with phenotypic BMI (p<2 × 10^−16^) and both predictors accounted for 18% of variation in phenotypic BMI.

## Epigenetic BMI score is associated with self-reported disease history

Table 2 presents results of logistic regressions carried out to test whether epigenetic BMI score is associated with self-reported disease history. Epigenetic BMI was positively associated with type 2 diabetes (OR 2.25, 95 % CI 1.74, 2.94), cardiovascular disease (OR 1.44, 95% CI 1.23, 1.69), and high blood pressure (OR 1.21, 95% CI 1.13, 1.48), all at p<0.0011 after Bonferroni correction. Phenotypic BMI was associated with the same self-reported disease variables: type 2 diabetes (OR 1.83, 95 % CI 1.45, 2.30), cardio vascular disease (OR 1.30, 95 % CI 1.11, 1.51) and high blood pressure (OR 1.57, 95% CI 1.36, 1.82). An additive model with both epigenetic and phenotypic BMI scores as predictors was associated with type 2 diabetes and cardiovascular disease.

**Table 2:** Logistic regression of epigenetic BMI score on self-reported disease history.

| Epigenetic BMI |  |  |  |  | Phenotypic BMI |  |  |  | Additive model |  |  |  |  |  |  |
| --- | --- | --- | --- | --- | --- | --- | --- | --- | --- | --- | --- | --- | --- | --- | --- |
|  |  |  |  |  |  |  |  |  | Epigenetic BMI |  |  | Phenotypic BMI |  |  |  |
|  | n | standard<br>ised<br>beta | OR (95%<br>CI) | raw p | n | standard<br>ised<br>beta | OR (95%<br>CI) | raw p | n | standar<br>dised<br>beta | OR (95%<br>CI) | raw p | standardi<br>sed beta | OR (95%<br>CI) | raw p |
| High blood pressure | 901 | 0.257 | 1.21 (1.13, 1.48) | 2.4x10<sup>-04</sup> | 902 | 0.450 | 1.57 (1.36, 1.82) | 9.8x10<sup>-10</sup> | 900 | 0.134 | 1.14 (0.99, 1.32) | 0.071 | 0.410 | 1.51 (1.30, 1.76) | 9.7x10<sup>-08</sup> |
| Type 1 diabetes | 901 | 0.827 | 2.29 (1.32, 3.94) | 0.003 | 902 | 0.642 | 1.90 (1.16, 2.98) | 0.007 | 900 | 0.684 | 1.98 (1.09, 3.55) | 0.022 | 0.436 | 1.55 (0.90, 2.51) | 0.090 |
| Type 2 diabetes | 901 | 0.811 | 2.25 (1.74, 2.94) | 1.4x10<sup>-09</sup> | 902 | 0.604 | 1.83 (1.45, 2.30) | 2.4x10<sup>-07</sup> | 900 | 0.690 | 1.99 (1.51, 2.64) | 1.1x10<sup>-06</sup> | 0.419 | 1.52 (1.19, 1.94) | 0.001 |
| High cholesterol | 900 | 0.071 | 1.07 (0.93, 1.23) | 0.318 | 901 | 0.145 | 1.16 (1.01, 1.33) | 0.039 | 899 | 0.029 | 1.03 (0.89, 1.19) | 0.694 | 0.139 | 1.15 (0.99, 1.33) | 0.061 |
| Cardiovascular disease | 901 | 0.363 | 1.44 (1.23, 1.69) | 5.8x10<sup>-06</sup> | 902 | 0.260 | 1.30 (1.11, 1.51) | 7.6x10<sup>-04</sup> | 900 | 0.317 | 1.37 (1.17, 1.62) | 1.7x10<sup>-04</sup> | 0.164 | 1.18 (1.00, 1.38) | 0.044 |
| Leg pain | 901 | 0.045 | 0.51 (0.91, 1.20) | 0.511 | 902 | 0.079 | 1.08 (0.95, 1.24) | 0.250 | 900 | 0.021 | 1.02 (0.89, 1.18) | 0.776 | 0.072 | 1.07 (0.93, 1.24) | 0.317 |
| Blood circulation | 899 | -0.241 | 0.79 (0.65, 0.95) | 0.013 | 900 | -0.002 | 0.10 (0.83, 1.19) | 0.986 | 898 | -0.267 | 0.77 (0.62, 0.93) | 0.009 | 0.075 | 1.08 (0.89, 1.31) | 0.446 |
| Stroke | 901 | 0.245 | 1.28 (0.95, 1.72) | 0.105 | 902 | -0.032 | 0.97 (0.71, 1.30) | 0.834 | 900 | 0.278 | 1.32 (0.97, 1.80) | 0.079 | -0.116 | 0.89 (0.64, 1.21) | 0.475 |
| Neoplasm | 901 | 0.050 | 1.05 (0.86, 1.28) | 0.627 | 902 | 0.116 | 1.12 (0.92, 1.35) | 0.234 | 900 | 0.012 | 1.01 (0.82, 1.05) | 0.913 | 0.112 | 1.12 (0.91, 1.36) | 0.275 |
| Thyroid | 900 | 0.011 | 1.01 (0.79, 1.29) | 0.931 | 901 | 0.111 | 1.12 (0.89, 1.39) | 0.327 | 899 | -0.030 | 0.97 (0.75, 1.26) | 0.823 | 0.119 | 1.13 (0.89, 1.42) | 0.318 |
| Parkinson's disease | 901 | -0.298 | 0.74 (0.29, 1.04) | 0.519 | 902 | -0.361 | 0.70 (0.24, 1.67) | 0.471 | 900 | -0.213 | 0.81 (0.31, 2.03) | 0.661 | -0.294 | 0.75 (0.25, 1.89) | 0.575 |
| Arthritis | 900 | -0.094 | 0.91 (0.79, 1.04) | 0.173 | 901 | 0.140 | 1.15 (1.03, 1.32) | 0.042 | 899 | -0.157 | 0.86 (0.74, 0.99) | 0.032 | 0.191 | 1.21 (1.05, 1.40) | 0.009 |
| Allergy | 901 | -0.002 | 0.10 (0.86, 1.15) | 0.974 | 902 | 0.077 | 1.08 (0.94, 1.24) | 0.280 | 900 | -0.032 | 0.97 (0.83, 1.13) | 0.683 | 0.087 | 1.09 (0.94, 1.27) | 0.249 |

## DISCUSSION

Using DNA methylation data, we constructed an epigenetic score for BMI to test its association with health, cognitive and lifestyle outcomes in an independent dataset, the LBC1936. We found that epigenetic BMI score is associated with poorer physical health (time taken to walk 6 metres, lung function and general physical health), higher levels of HbA1c, blood derived C-reactive protein, triglycerides and HDL cholesterol (all biomarkers for metabolic syndrome), poorer crystallised intelligence, lower physical health-related quality of life, a lower self-reported physical activity and greater social deprivation. Our analysis also showed that epigenetic BMI score is associated with self-reported type 2 diabetes (a finding previously reported by Wahl et al., who identified diabetes via clinical diagnosis, or HbA1c ≥ 6.5% (3)), cardiovascular disease and high blood pressure: health outcomes for which obesity is a major risk factor.

In almost all analyses, phenotypic BMI accounted for a greater proportion of variance of outcome variables than epigenetic BMI, but phenotypic and epigenetic BMI scores together accounted for a greater proportion of variance still. This additive effect suggests that epigenetic and phenotypic BMI scores account for at least partially independent proportions of variance. It is possible that some these partially independent associations are driven by correlations between the CpG probes used to create the epigenetic BMI score and the outcome variables of interest. This could mean that our epigenetic score for BMI is capturing the downstream effects of obesity, such as biological processes associated with poor metabolic health, which phenotypic BMI (a more ‘blunt’ measure of metabolic health) does not account for directly.

We found that an epigenetic score for BMI accounted for 10% of the variance in phenotypic BMI, a higher proportion than that reported by Shah et al. (6), whose epigenetic BMI score accounted for 6.9% of the variance in phenotypic BMI in the LBC1936. This improvement could be due to our larger discovery sample (5,200 vs 1,366), and/or because weights for our epigenetic BMI score were calculated using a single regression model, with BMI residuals as the outcome and CpGs as predictors, rather than separate models for each CpG predictor (see Appendix 1). Shah et al also found that an epigenetic BMI score improved phenotype prediction when combined with a genetic score for BMI from 8% to 14% (6). By including both an epigenetic and a polygenic score for BMI as predictors in our linear regression model, we improved the proportion of variance in phenotypic BMI explained from 10% to 18%. However, our polygenic score for BMI did not associate with any variables when included alongside epigenetic BMI score and phenotypic BMI as predictors in regression models. In the majority of these models, polygenic BMI score had little effect; associations between epigenetic BMI score and outcome variables remained, particularly for physical health and biomarker variables. It could be the case that the epigenetic BMI score encompasses the interaction between the polygenic BMI score and outcome variables implicated in biological processes related to obesity (as opposed to cognitive or lifestyle variables). If this is the case, there appears to be little benefit in the inclusion of a polygenic score for BMI alongside epigenetic BMI and phenotypic BMI scores in assessing association with biological and physical health-related variables.

Our results demonstrate that BMI (and epigenetic BMI) is associated with obesity-related adverse health and thus, is a clinically-relevant metric for measuring body mass. However, the BMI has been criticised due to its assumption that a low body mass equates to metabolic health. Dvorak et al. (16) found that BMI failed to distinguish between healthy individuals and individuals who were metabolically obese, but of normal weight despite differences in total fat mass, body fat percentage, subcutaneous fat and visceral abdominal adiposity. It has also been shown that waist to height ratio (WHtR) and waist/height^0.5^ (WHT.5R) are better predictors of whole body fat percentage and visceral adipose tissue mass than BMI (17). Despite this, the BMI continues to be widely used in clinical settings. A development upon this study would be to test whether an epigenetic score based on WHtR or WHT.5R can further increase the proportion of variance in obesity-related outcomes explained by epigenetic BMI.

A limitation of this study is the inability to make strong directional inferences; it is impossible to tell whether differential DNA methylation is a mediator of the causal disease pathway, or occurs in response to disease processes. Recent progress has been made in this area using a Mendelian Randomisation approach (18), which takes SNPs as a proxy for both the methylation profile and outcome of interest, in order to gain insight into the direction of causality. Longitudinal studies are another important method for gaining insight into the role of DNA methylation in the pathophysiology of obesity-related disease. Several studies have analysed DNA methylation from umbilical cord blood as a baseline measure for longitudinal epigenetic data. This is then analysed this in tandem with longitudinal data on health-related outcomes in order to capture epigenetic changes that may predate disease onset (for review see 19). These cohorts must have deep phenotyping of lifestyle and environmental factors (including prenatal factors such as maternal diet and smoking) that may confound epigenetic changes. Another developing line of research is investigating epigenetic changes associated with weight-loss interventions such as diet programmes or gastric bypass surgery (20, 21).

A recent study by Richmond et al. (22) combined several of these methods to detect causality between BMI and methylation at *HIF3A* (a gene linked to metabolism and obesity, and associated with adiposity in multiple EWAS (23, 24)), including bidirectional Mendelian randomisation analysis, longitudinal analysis of *HIF3A* methylation, and analysis of association between maternal pregnancy BMI and *HIF3A* methylation in offspring at birth (using cord blood). Results across each method indicated that a higher BMI causally influenced higher *HIF3A* methylation, suggesting that alterations in DNA methylation are a downstream effect of obesity. Further longitudinal studies and studies of naturalistic interventions are needed to elucidate the direction of causal association between obesity and DNA methylation, and to provide mechanistic insight into the pathophysiology of obesity-related disease.

We have demonstrated that an epigenetic score for BMI is associated with poorer physical health and health-related quality of life, biomarkers for metabolic syndrome and major diseases for which obesity is a major risk factor, social deprivation, physical inactivity and lower crystallised intelligence. An epigenetic score for BMI based on DNA methylation increased the amount of variance in health outcomes accounted for by phenotypic BMI score alone, demonstrating the value in using epigenetic information for constructing more accurate risk prediction for obesity-related biomarkers, health conditions and disease.

## ACKNOWLEDGEMENTS

The authors thank all LBC1936 study participants and research team members who have contributed, and continue to contribute, to ongoing LBC1936 study. The Lothian Birth Cohort 1936 is supported by Age UK (Disconnected Mind programme) and the Medical Research Council [MR/M01311/1]. Methylation typing was supported by Centre for Cognitive Ageing and Cognitive Epidemiology (Pilot Fund award), Age UK, The Wellcome Trust Institutional Strategic Support Fund, The University of Edinburgh, and The University of Queensland. This work was conducted in the Centre for Cognitive Ageing and Cognitive Epidemiology, which is supported by the Medical Research Council and Biotechnology and Biological Sciences Research Council [MR/K026992/1], and which supports Ian Deary. Generation Scotland received core support from the Chief Scientist Office of the Scottish Government Health Directorates [CZD/16/6] and the Scottish Funding Council [HR03006]. Genotyping of the GS:SFHS samples was carried out by the Genetics Core Laboratory at the Wellcome Trust Clinical Research Facility, Edinburgh, Scotland and was funded by the Medical Research Council UK and the Wellcome Trust (Wellcome Trust Strategic Award “STratifying Resilience and Depression Longitudinally” [(STRADL) Reference 104036/Z/14/Z].

